# 3D mechanical confinement directs muscle stem cell fate and function

**DOI:** 10.1101/2024.10.03.616478

**Authors:** GaYoung Park, Josh A. Grey, Foteini Mourkioti, Woojin M. Han

## Abstract

Muscle stem cells (MuSCs) play a crucial role in skeletal muscle regeneration, residing in a niche that undergoes dimensional and mechanical changes throughout the regeneration process. This study investigates how three-dimensional (3D) confinement and stiffness encountered by MuSCs during the later stages of regeneration regulate their function, including stemness, activation, proliferation, and differentiation. We engineered an asymmetric 3D hydrogel bilayer platform with tunable physical constraints to mimic the regenerating MuSC niche. Our results demonstrate that increased 3D confinement maintains *Pax7* expression, reduces MuSC activation and proliferation, inhibits differentiation, and is associated with smaller nuclear size and decreased H4K16ac levels, suggesting that mechanical confinement modulates both nuclear architecture and epigenetic regulation. MuSCs in unconfined two-dimensional (2D) environments exhibited larger nuclei and higher H4K16ac expression compared to those in more confined 3D conditions, leading to progressive activation, expansion, and myogenic commitment. This study highlights the importance of 3D mechanical cues in MuSC fate regulation, with 3D confinement acting as a mechanical brake on myogenic commitment, offering novel insights into the mechano-epigenetic mechanisms that govern MuSC behavior during muscle regeneration.

## INTRODUCTION

Stem cells respond to mechanical cues derived from their microenvironment, which regulate cellular processes such as proliferation, differentiation, and self-renewal (*1–3*). In two-dimensional (2D) systems, cells freely spread, sense the underlying stiffness, and regulate their behavior. In three-dimensional (3D) environments, cells are physically confined by the surrounding matrix, which exerts mechanical forces that regulate cellular and nuclear volume and influence their function in ways not seen in 2D systems (*3–10*). Physical confinement – dictated by ECM pore size, viscoelasticity, and degradability – introduces additional complexity that modulates stem cell biology (*3*). While 2D stem cell mechanobiology studies have revealed considerable insights into how matrix stiffness regulates stem cell function (*11*), they overlook key 3D-specific cues, such as confinement and asymmetric mechanical properties, which are crucial for understanding how stem cells function in health and disease.

Muscle stem cells (MuSCs) navigate both 2D and 3D environmental contexts due to the unique geometry and properties of their niche, which dynamically change during regeneration. In homeostasis, this niche is asymmetrically 3D, with polarized physicochemical cues originating from the basement membrane on one side and the myofiber on the other, creating a distinct microenvironment that regulates MuSC function (*12*, *13*). Following muscle injury, the necrotic myofiber is removed, leaving behind the basement membrane, often called the “ghost fiber,” which acts as a 2D-like scaffold (*14*, *15*). MuSCs use this template to expand and generate highly proliferative myogenic cells that eventually fuse in a density-dependent manner to form regenerating myofibers (*14*, *15*). As regenerating myofibers emerge, MuSCs gradually decrease proliferation, self-renew, exit the cell cycle, and return to quiescence, a process critical for maintaining their long-term regenerative capacity (*15*, *16*). During this transition, MuSCs re-enter the asymmetric 3D environment, where they are confined and compressed by the niche; however, whether and how these extrinsic physical cues regulate MuSC behavior— particularly their slowed proliferation, self-renewal, and quiescence re-entry—remains unknown.

The MuSC niche is mechanically dynamic, characterized by forces including compression, tension, ECM stiffness, intramuscular pressure, and confinement (*17*, *18*). Among these, stiffness and compression are critical in regulating MuSC self-renewal and quiescence (*19–24*). For instance, studies using muscle-mimetic 2D hydrogels with 12 kPa stiffness have shown that MuSCs expand under such conditions (*19*, *20*, *22*), while higher stiffnesses lead to persistent activation (*21*). However, 2D studies fail to capture the complexity of the 3D mechanical environment that MuSCs encounter during later stages of regeneration, particularly the effects of asymmetric stiffness and confinement. Recent findings suggest that static compression promotes MuSC quiescence (*24*), further highlighting the role of other mechanical cues that direct MuSC function. As regenerating muscle undergoes dynamic biomechanical changes (*21*, *25*), diverse mechanical cues likely influence MuSC function, including proliferation, self-renewal, and their return to quiescence. Yet, the specific roles of 3D mechanical factors like confinement and asymmetric stiffness in regulating MuSC behavior remain unexplored, largely due to the inherent challenges of studying these factors *in vivo*.

In this study, we engineered and applied a 3D asymmetric bilayer hydrogel platform that mimics distinct stages MuSC niche during muscle regeneration and found that 3D mechanical confinement promotes the maintenance of *Pax7* expression of MuSCs, delays their activation and differentiation, and restricts nuclear size in a stiffness and confinement-dependent manner. Notably, such physical constraints reduced H4K16ac levels, a marker of chromatin accessibility and myogenic gene activation, suggesting that mechanical confinement regulates MuSC function through mechano-epigenetic mechanisms. These findings provide new insights into how 3D confinement acts as a mechanical brake that restricts continued MuSC expansion and myogenic commitment, signaling MuSCs toward niche repopulation.

## RESULTS

### Facile assembly of bilayer synthetic hydrogel system that mimics asymmetrically 3D regenerating MuSC niche

The MuSC niche exhibits dynamic asymmetry and confinement across unconfined 2D and confining 3D contexts during regeneration (**Fig. 1A**). Confinement refers to the restriction of MuSCs within a controlled 3D space, limiting their ability to expand and interact with their surroundings, while stiffness modulates the degree of physical resistance the cells experience (*3*). To investigate the effects of confinement on MuSC function, we engineered an asymmetric 3D hydrogel platform using well-established norbornene-terminated multi-arm poly(ethylene) glycol (PEG-NB) to create a biofunctional hydrogel bilayer, with cells seeded within the hydrogel interface (**Fig. 1B**). By adjusting the crosslink density, we can tune the stiffness of each hydrogel layer to match the physiologically relevant Young’s modulus range for regenerating skeletal muscle (5-35 kPa) (**Fig. 1C**) (*21*).

**Figure 1.**
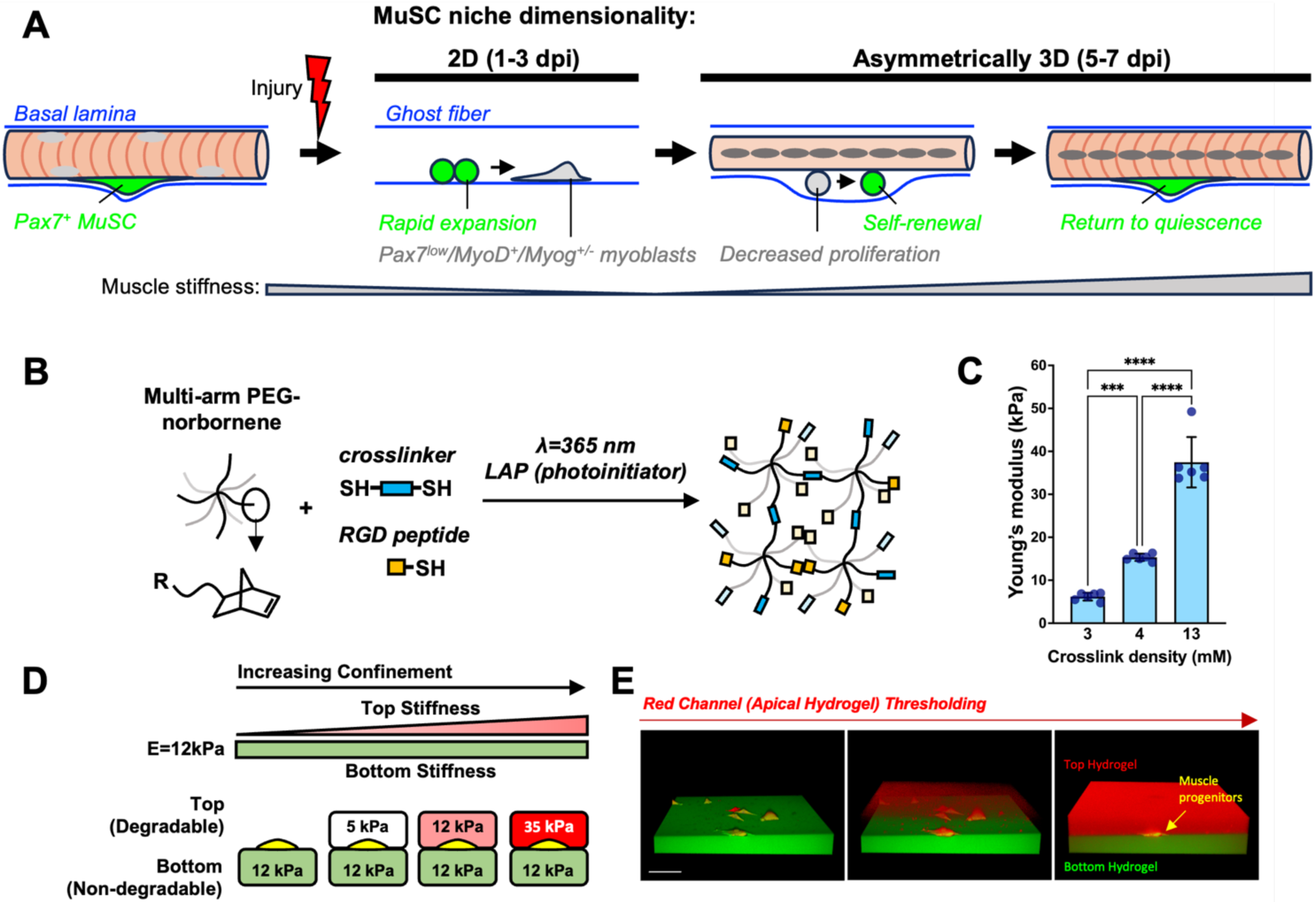
Facile assembly of bilayer hydrogel system that mimics asymmetrically 3D regenerating MuSC niche. **(A)** Quiescent MuSCs reside between the basal lamina and the myofiber under normal conditions. Upon injury, the myofiber undergoes necrosis, and activated MuSCs utilize the remaining basal lamina, or “ghost fiber,” as a 2D scaffold to rapidly expand and differentiate. As regenerating myofibers emerge, the MuSCs are compressed, leading to decreased proliferation, increased self-renewal, and eventual return to quiescence. **(B)** Schematic of multi-arm PEG-norbornene hydrogel. Synthetic hydrogel can be functionalized with cell-adhesive ligands and thiol-flanked crosslinkers using light. **(C)** Young’s modulus of hydrogel as a function of crosslink density. n=6 hydrogels. *** p<0.001, **** p<0.0001 via 1-way ANOVA with Tukey’s post-hoc tests. **(D)** Schematic of bilayer hydrogel conditions: 2D mimics the ghost fiber environment; 5/12 kPa represents a soft regenerating asymmetric niche; 12/12 kPa represents a homeostatic asymmetric niche; 35/12 kPa represents a stiff regenerating niche. Stiffer hydrogels provide greater confinement. **(E)** 3D-reconstructed confocal micrograph demonstrating sandwiched C2C12 myoblasts between two hydrogel layers. Scale bar: 100 µm.

To create this biofunctional hydrogel bilayer, we first seeded C2C12 myoblasts on a flat, non-degradable RGD-functionalized 12 kPa hydrogel, which has previously been shown to support rapid expansion of self-renewing MuSCs (*19*, *20*, *22*). After allowing 16 hours for cell adhesion, we cast and polymerized a second layer of protease-degradable RGD-functionalized hydrogel with variable stiffness—5 kPa (soft), 12 kPa (intermediate), and 35 kPa (stiff)—on top of the attached cells (**Fig. 1D**). Here, 2D mimics the unconfined ghost fiber environment; 5/12 kPa represents a soft regenerating asymmetric niche; 12/12 kPa represents a homeostatic asymmetric niche; 35/12 kPa represents a stiff regenerating niche (**Fig. 1A**) (*21*), where degree of confinement increases with stiffness (*3*). To ensure a stable hydrogel interface, we carefully controlled the time before polymerizing the second hydrogel layer upon deposition. This approach promotes polymer entanglements through polymer diffusion into the bottom hydrogel layer pre-polymerization (*26–28*), resulting in a highly reproducible method for sandwiching muscle cells between two uniform hydrogel layers with distinct mechanical properties (**Fig. 1E**). Thus, we successfully devised an asymmetric 3D hydrogel platform that allows for the investigation of 3D mechanical cues on MuSC function.

### Dynamic Pax7EGFP reporter MuSCs enable longitudinal tracking of Pax7 expression ex vivo

The *Pax7* transcription factor is the characteristic marker of MuSCs in adult skeletal muscle, whereas activated and differentiating MuSCs gradually lose *Pax7* expression (*29–31*). To facilitate longitudinal monitoring of dynamic *Pax7* expression ex vivo, we used primary MuSCs isolated from the hindlimb muscles of previously described Pax7EGFP reporter mice (*32*), where EGFP is directly driven by *Pax7* expression. We first tested whether the Pax7EGFP reporter could be used to distinguish the temporally dynamic *Pax7* expression levels of MuSCs on a laminin/collagen-coated tissue culture plastic and 12 kPa RGD-functionalized 2D hydrogel, where compliant 12 kPa substrates promote the expansion of *Pax7*+ MuSCs while MuSCs on tissue culture plastics rapidly lose their *Pax7* expression (*19*). Freshly isolated MuSCs exhibited bright and intense Pax7EGFP signals in both conditions (**Supp. Fig. 1A**). As expected, MuSCs on both substrates gradually lost the Pax7EGFP signal over the culture duration, but MuSCs maintained on the 12 kPa hydrogels exhibited significantly higher mean Pax7EGFP intensity over time, particularly by 120 hours (**Supp. Fig. 1B-D**). In addition, there was a significantly larger fraction of MuSCs exhibiting elevated Pax7EGFP intensity on the 12 kPa hydrogels compared to the plastic substrates (**Supp. Fig. 1E, F**). These results demonstrate the feasibility of using primary Pax7EGFP MuSCs to longitudinally monitor the temporal kinetics of *Pax7* expression levels on engineered hydrogel systems.

### 3D confinement limits MuSC proliferation and sustains *Pax7* expression

Physical and mechanical stimuli dictate cellular behavior of how stem cells sense and respond to their local microenvironment (*3*). Thus, we next introduced primary Pax7EGFP MuSCs in the 3D bilayer and on 2D hydrogels to investigate how confinement regulates MuSC expansion. By longitudinally tracking the cells over time, we observed that MuSCs maintained on the 2D 12 kPa hydrogels expanded rapidly over 5 days (**Fig. 2A, B**), consistent with previous observations showing that soft substrates promotes MuSC expansive growth (*19*, *20*, *22*). In contrast, we found that MuSCs maintained in 3D bilayer hydrogels exhibited significantly decreased expansion compared to the 2D condition (**Fig. 2A, B**). In addition, we observed that MuSCs in 3D bilayer hydrogels slowly expanded in isolated clusters, where the colony density was the lowest in the stiffest 3D (35 kPa/12 kPa) condition (**Fig. 2A, C**), suggesting that various degree of physical constraints differentially influence in MuSC expansion.

**Figure 2.**
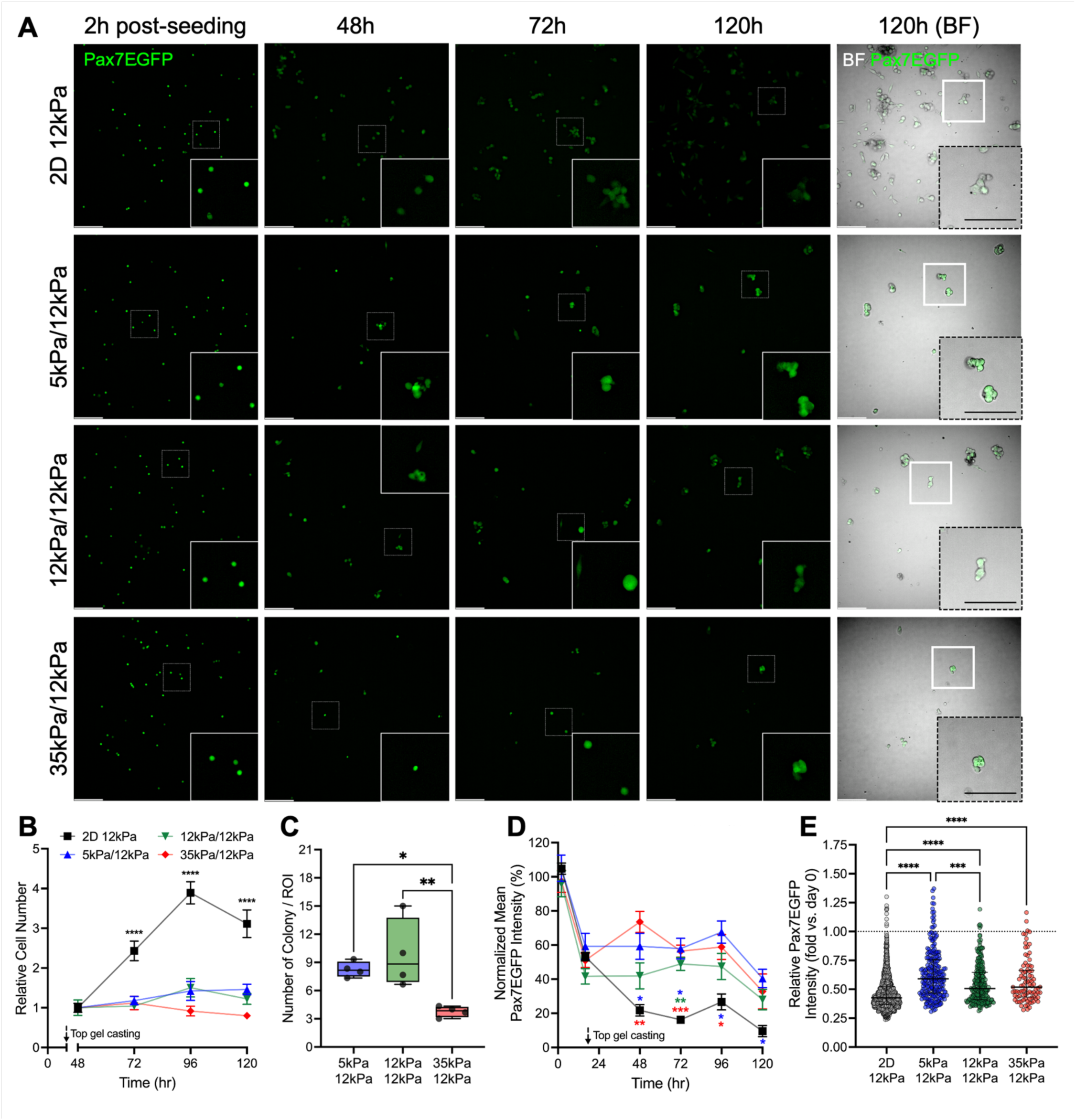
3D confinement limits MuSC proliferation and myogenic commitment. **(A)** Representative micrographs of Pax7EGFP MuSCs cultured on 2D 12 kPa hydrogel and 3D bilayer hydrogels of varying stiffnesses. Scale bar: 100 µm. BF: Bright Field. **(B)** Relative MuSC number within 2D and 3D hydrogels. n=4 hydrogels. **** p<0.0001 vs all 3D groups via two-way ANOVA with Dunn’s post-hoc. **(C)** Number of MuSC colonies per region-of-interest (ROI). n=4 hydrogels. * p<0.05, ** p<0.01 via one-way ANOVA with Tukey’s post-hoc. **(D)** Normalized Pax7EGFP intensity of MuSCs over time. n=4 hydrogels. * p<0.05, ** p<0.01, *** p<0.001 via two-way ANOVA with Dunn’s post-hoc. **(E)** Relative Pax7EGFP intensity of MuSCs on day 5 normalized to day 0 intensity. n=88-1451 cells analyzed. *** p<0.001, **** p<0.0001 via Kruskal-Wallis with Dunn’s post-hoc.

To investigate whether confinement can also regulate *Pax7* expression, we next quantified Pax7EGFP intensity at multiple time points over 5 days. Freshly isolated MuSCs exhibit bright Pax7EGFP intensity 2 hours post-seeding, which rapidly declines by 16 hours, just before the second hydrogel casting (**Fig. 2A, D**). As isolated MuSCs are known to spontaneously activate (*33*), this decrease in intensity further corroborates the validity of our system. MuSCs maintained on the 2D hydrogels continue to lose Pax7EGFP intensity over time (**Fig. 2A and D**), indicating progressive activation and a loss of stemness. Interestingly, MuSCs maintained in 3D bilayer hydrogels sustained higher Pax7EGFP intensity, with levels significantly greater than those observed in the 2D condition (**Fig. 2A, D**). Notably, a significantly higher fraction of MuSCs maintained in 3D bilayer hydrogels exhibited high Pax7EGFP intensity compared to those on 2D hydrogels by day 5 (**Fig. 2A, E**), indicating sustained *Pax7* expression. Performing PAX7 immunostaining on Pax7EGFP MuSCs on day 5 in the 3D soft (5 kPa/12 kPa) and 2D conditions, which showed the highest and lowest Pax7EGFP intensities, respectively, revealed no differences in PAX7 positivity (**Supp. Fig. 2A, B**). This suggests that while most Pax7EGFP+ cells express comparable PAX7 protein in both 2D and 3D contexts, the reduction in Pax7EGFP intensity in 2D hydrogels likely reflects decreased Pax7 transcript levels, pointing to a further activation state that is not captured by immunostaining alone, which only provides an on/off readout of PAX7 expression.

### 3D confinement delays MuSC activation

MuSC activation is a critical step in transitioning from quiescence to myogenic commitment following disruption of stem cell niche, signaling the start of myogenesis (*14*, *15*). Shortly after early activation events, including the loss of cellular projections (*34*, *35*) and FOS induction (*36*), activated MuSCs robustly express the myogenic regulatory factor MyoD (*37*). To assess how 3D confinement impacts MuSC activation, we cultured Pax7EGFP MuSCs within the 3D bilayer and on 2D hydrogels for 3 and 5 days, then evaluated the activation marker MyoD. By day 3, approximately 70% of Pax7EGFP+ MuSCs on 2D hydrogels expressed MyoD, indicating spontaneous activation (**Fig. 3A, B**). In contrast, MyoD positivity was significantly reduced to about 40% in Pax7EGFP+ MuSCs cultured in 3D bilayer hydrogels compared to the 2D condition (**Fig. 3A, B**). By day 5, approximately 80% of Pax7EGFP+ MuSCs on 2D hydrogels expressed MyoD, while Pax7EGFP+ MuSCs in 3D bilayer hydrogels continued to exhibit significantly fewer MyoD-positive nuclei (**Fig. 3C, D**), further demonstrating the effect of confinement in influencing the activation status of MuSCs. Interestingly, the most confining and stiff 3D conditions further suppressed MyoD expression compared to the soft and intermediate 3D conditions (**Fig. 3C, D**). These results collectively indicate that 3D confinement delays MuSC activation in a confinement and stiffness-dependent manner.

**Figure 3.**
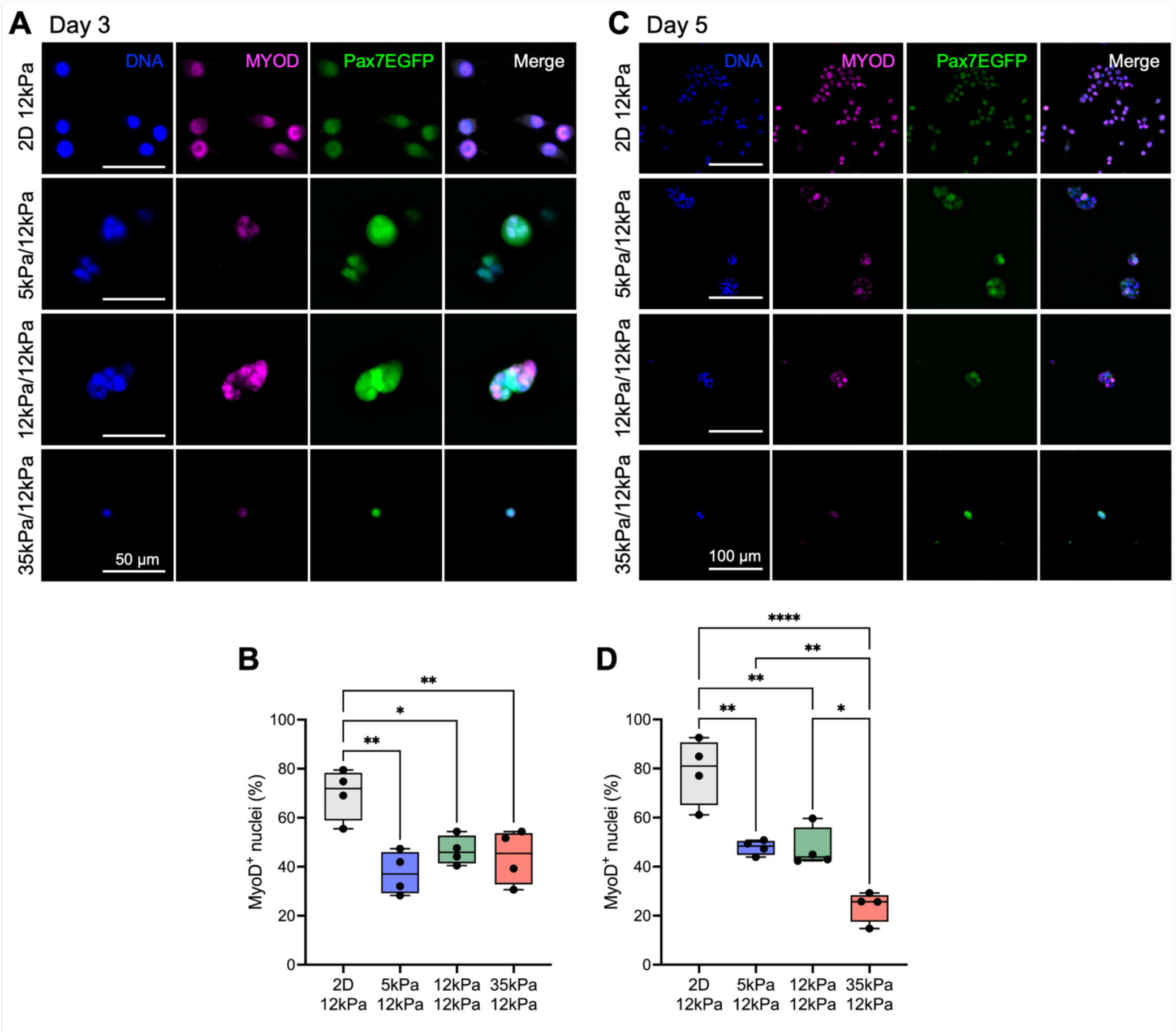
3D confinement delays MuSC activation in a confinement and stiffness-dependent manner. **(A)** MyoD staining on day 3. Scale bar: 50 µm. **(B)** Percentage of MyoD+ nuclei on day 3. n=4 hydrogels. * p<0.05, ** p<0.01 via one-way ANOVA with Tukey’s post-hoc. **(C)** MyoD staining on day 5. Scale bar: 100 µm. **(D)** Percentage of MyoD+ nuclei on day 5. n=4 hydrogels. * p<0.05, ** p<0.01, **** p<0.0001 via one-way ANOVA with Tukey’s post-hoc.

### 3D confinement inhibits MuSC differentiation

Myogenin (MYOG) expression marks the beginning of differentiation, promoting the maturation of activated MuSCs/myoblasts into fusion-competent myocytes (*37*). To assess whether confinement delays MuSC differentiation onset, we immunostained MuSCs with MYOG. Approximately 35% of MuSCs cultured on the 2D hydrogels expressed MYOG by day 5, while MuSCs maintained in 3D bilayer hydrogels exhibited significantly fewer MYOG+ nuclei compared to the 2D condition (**Fig. 4A, B**). Notably, MuSCs in the stiffest 3D bilayer hydrogel showed no MYOG+ nuclei (**Fig. 4A, B**), suggesting that confinement delays the onset of myogenic differentiation, particularly under more confining and stiff 3D conditions.

**Figure 4.**
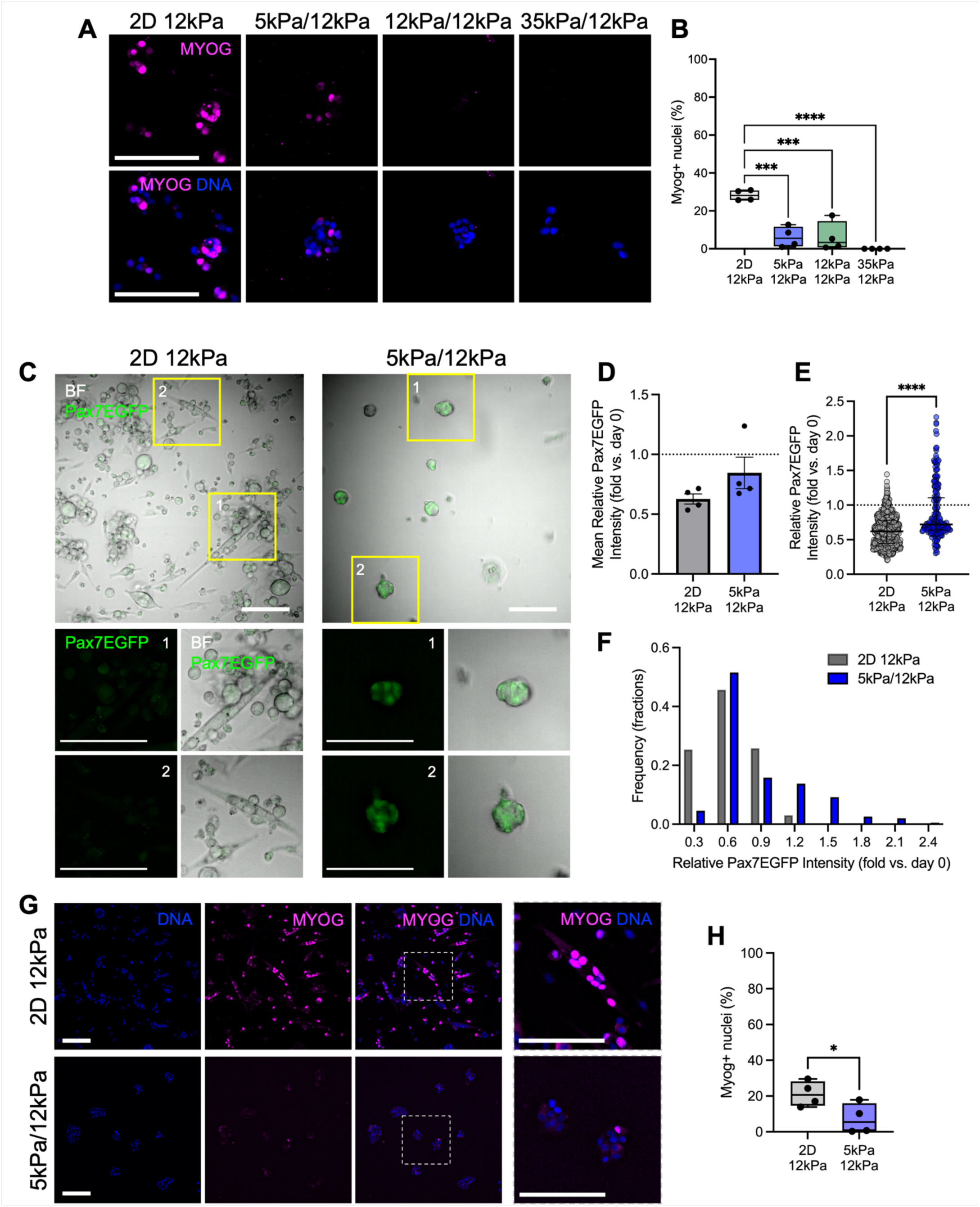
3D confinement inhibits MuSC differentiation. **(A)** Myogenin staining on day 5. Scale bar: 100 µm. **(B)** Percentage of myogenin+ nuclei on day 5. n=4 hydrogels. *** p<0.001, **** p<0.0001 via one-way ANOVA with Tukey’s post-hoc. **(C)** Representative micrographs of Pax7EGFP MuSCs cultured on 2D 12 kPa hydrogel and 3D 5/12 kPa soft bilayer hydrogel on day 7. Scale bar: 100 µm. BF: Bright Field. **(D)** Mean relative MuSC number within 2D and 3D hydrogels. n=4 hydrogels. **(E)** Relative Pax7EGFP intensity of MuSCs on day 5 normalized to day 0 intensity. n=196-725 cells analyzed. **** p<0.0001 via two-tailed Mann-Whitney U test. **(F)** Frequency distribution of relative Pax7EGFP intensity of MuSCs normalized to day 0 intensity. **(G)** Myogenin staining on day 7. Scale bar: 100 µm. (**H)** Percentage of myogenin+ nuclei on day 7. n=4 hydrogels. * p<0.05 via two-tailed t-test.

To further explore the effects of 3D confinement on MuSC differentiation, we extended the culture period to 7 days, focusing on the 2D and 3D soft (5 kPa/12 kPa) conditions, which showed the largest differences in Pax7EGP intensity on day 5 (**Fig. 2D, E**). On unconfined 2D hydrogels, MuSCs proliferated to high cell density and spontaneously differentiated into large, elongated myotubes (**Fig. 4C**). In contrast, under 3D confining conditions, MuSCs remained undifferentiated and formed colony-like structures (**Fig. 4C**). In the 3D bilayer hydrogels, MuSCs exhibited significantly higher Pax7EGFP intensity compared to those on 2D hydrogels (**Fig. 4D-F**). Furthermore, colony-forming MuSCs in the 3D hydrogels had significantly fewer MYOG+ nuclei than those on the 2D hydrogels (**Fig. 4G, H**). Taken together, these results demonstrate that 3D confinement inhibits MuSC differentiation.

### 3D confinement restricts MuSC nuclei and colony size in a stiffness-dependent manner

Numerous studies in mechanobiology demonstrate that geometric constraints on a cell induce deformation, which in turn impacts nuclear and chromatin dynamics (*38*). To investigate nuclear alterations in MuSCs, we asked whether 3D confinement limits nuclear area and colony size. On day 3, the nuclear area decreased in a confinement-dependent manner, with the largest MuSC nuclear area observed on unconfined 2D hydrogels and the smallest in the stiffest, most confining 3D bilayer hydrogels (**Fig. 5A, B**). Additionally, we noted that larger nuclei exhibited more “nuclear puncta”, likely representing nucleoli and heterochromatin foci, which were absent in the stiffest 3D bilayer hydrogels (**Fig. 5A**). Within the 3D hydrogel conditions, the colony size – quantified by the number of nuclei – was also inversely correlated with increasing stiffness and confinement (**Fig. 5A, C**). This trend persisted through day 5, with both nuclear area and colony size remaining smaller in the more confined and stiffer 3D conditions to the 2D hydrogels (**Fig. 5D-F**), indicating that the level of physical constraint the cells experience is significantly affecting their nuclear size and number, and it is likely to impact their downstream cellular function.

**Figure 5.**
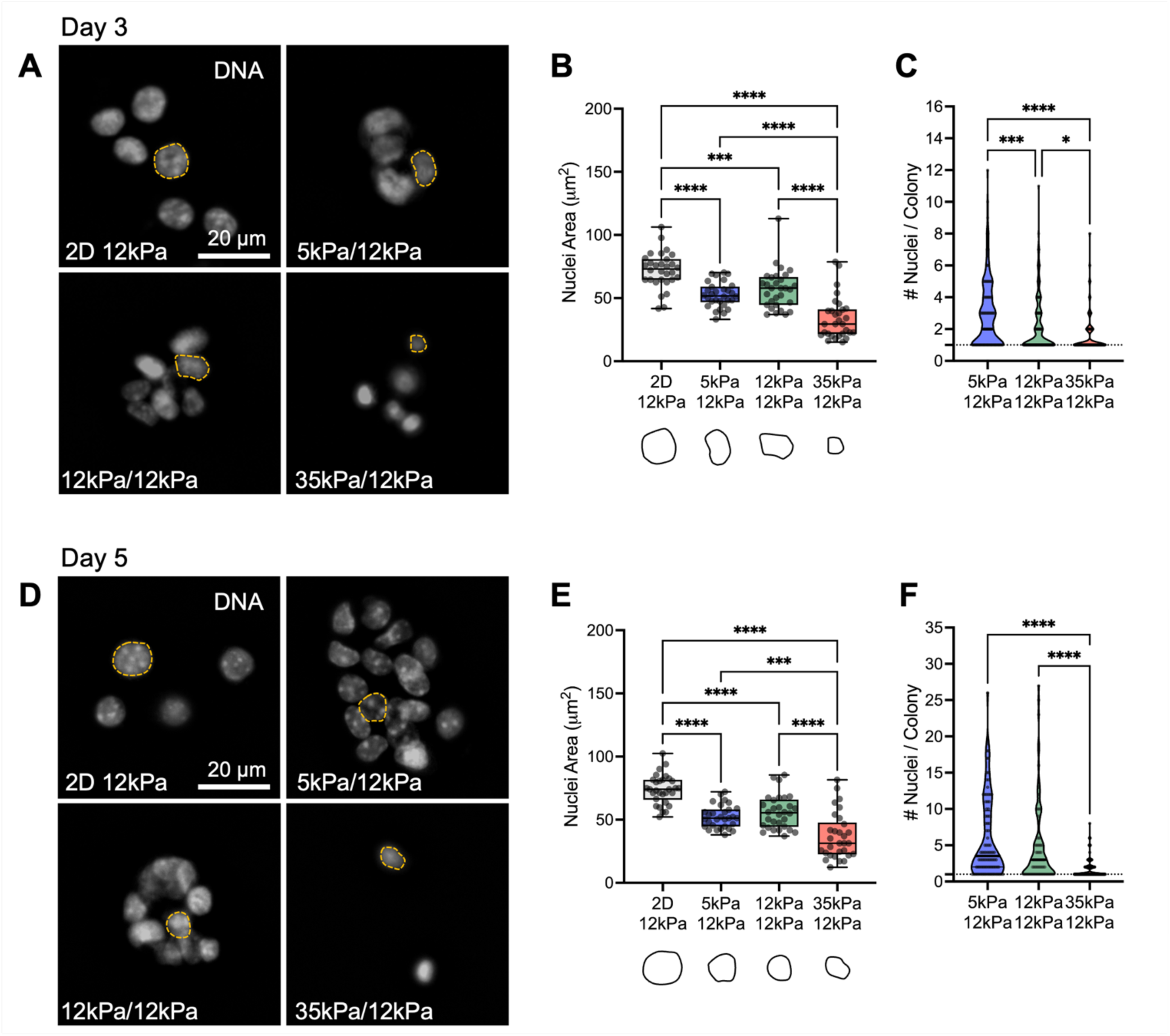
3D confinement restricts MuSC nuclei and colony size in a stiffness-dependent manner. **(A)** DNA (Hoechst) staining on day 3. Scale bar: 20 µm. **(B)** Quantification of nuclei area. n=30 nuclei analyzed. *** p<0.001, **** p<0.0001 via one-way ANOVA with Tukey’s post-hoc. **(C)** Quantification of number of nuclei per colony. n=148-166 colonies analyzed. * p<0.05, *** p<0.001, **** p<0.0001 via Kruskal-Wallis with Dunn’s post-hoc. **(D)** DNA (Hoechst) staining on day 5. Scale bar: 20 µm. **(E)** Quantification of nuclei area. *** p<0.001, **** p<0.0001 via one-way ANOVA with Tukey’s post-hoc. **(F)** Quantification of number of nuclei per colony. n=88-131 colonies analyzed. **** p<0.0001 via Kruskal-Wallis with Dunn’s post-hoc.

In this system, MuSCs control their confinement by deforming the surrounding environment and secreting proteases to degrade the top hydrogel. In the regenerating skeletal muscle, MuSCs secrete MMP2 and MMP9 to remodel basal lamina to expand and self-renew (*39*). To decouple these mechanisms in regulating MuSC nuclear area and colony size, we cultured MuSCs in soft and stiff 3D bilayer hydrogels, where the top hydrogels were either crosslinked with protease-degradable VPM (MMP2 and MMP9-sensitive)(*5*, *6*, *40*) or non-degradable PEG-dithiol. In the soft 3D bilayer hydrogels, MuSC nuclear size was significantly smaller in the non-degradable condition than in the protease-degradable condition (**Supp. Fig. 3A**), suggesting the matrix degradability contributes to the relief of confinement and allows the nuclei enlargement. There were no effects on the colony size, indicating that proliferation in soft 3D bilayer hydrogels is primarily governed by matrix stiffness rather than degradation. However, in the stiff 3D bilayer hydrogels, protease degradability had no effects on the nuclear area and colony size (**Supp. Fig. 3B**), suggesting that matrix stiffness dominates over any potential effect of matrix degradation in highly stiff environments. Stiffness likely imposes a mechanical limit on both nuclear size growth and colony expansion, regardless of matrix degradability. Therefore, in stiff environments, MuSCs are less able to overcome confinement through matrix degradation, and their behavior is primarily dictated by the stiffness and confinement of the surrounding environment.

### 3D confinement is associated with decreased H4K16 acetylation

Since we observed differences in the nuclear size as a function of confinement, we next investigated whether confinement regulates chromatin accessibility. Quiescent MuSCs possess highly condensed heterochromatin, where this condensed state is essential for preventing activation (*41*). Upon activation, MuSCs undergo nuclear enlargement, increase their glycolytic activity, and exhibit a significant increase in histone H4 lysine 16 acetylation (H4K16ac), marking a shift towards more accessible euchromatin and downstream activation of MyoD expression (*42–44*). On unconfined 2D hydrogels, nearly all MuSCs expressed H4K16ac+ nuclei by day 5 (**Fig. 6A, B**). In the confining 3D conditions, MuSCs expressed H4K16ac in a confinement-dependent manner, with the highest expression in soft 3D bilayer hydrogels and the lowest in stiff 3D bilayer hydrogels (**Fig. 6A, B**). Notably, MuSCs maintained in the stiff 3D (35 kPa/12 kPa) condition exhibited significantly fewer H4K16ac+ nuclei compared to both 2D unconfined and soft/intermediate 3D bilayer hydrogels (**Fig. 6A, B**). These findings suggest that 3D confinement not only limits nuclear size (**Fig. 5A-F**) but also suppresses chromatin accessibility marked by the reduction of H4K16ac levels. This effect is most pronounced in the stiffest 3D conditions, where MuSCs are less able to remodel their chromatin, likely impacting their transcriptional activation. Taken together, our results demonstrate that 3D confinement not only reduces MuSC activation and proliferation, but also restricts chromatin accessibility and gene regulation, suggesting potential epigenetic control mechanisms under 3D physical stress.

**Figure 6.**
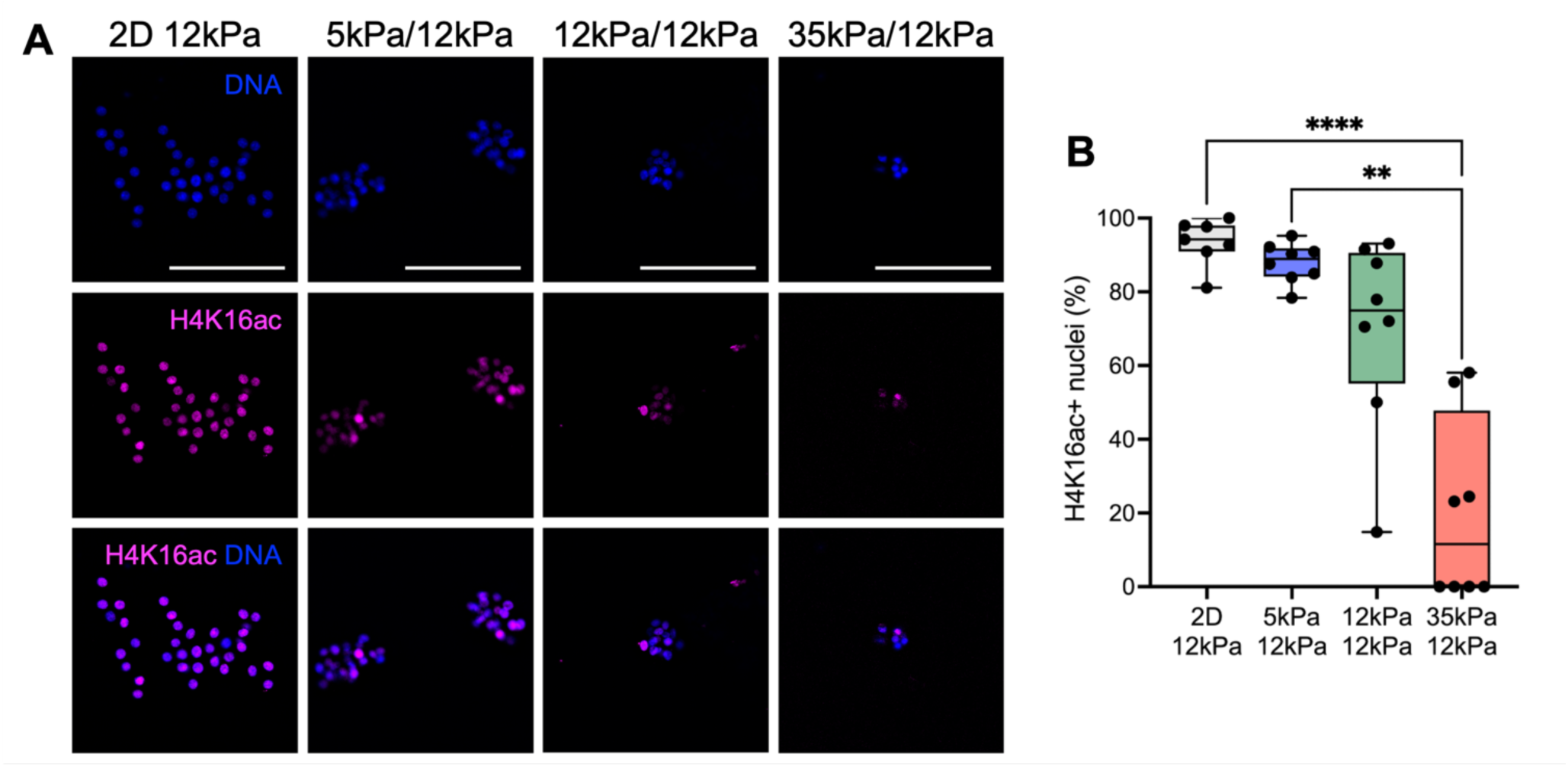
3D confinement is associated with decreased H4K16 acetylation. **(A)** H4K16ac staining on day 5. Scale bar: 100 µm. **(B)** Percentage of H4K16ac+ nuclei on day 5. n=7-8 hydrogels. ** p<0.01, **** p<0.0001 via Kruskal-Wallis with Dunn’s post-hoc.

## DISCUSSION

An unanswered question in muscle regenerative biology is how physical forces within the regenerating niche signal MuSCs to slow proliferation, self-renew, and return to quiescence as they re-enter the asymmetric, confining 3D environment, preventing chronic activation and overexpansion. During muscle regeneration, MuSCs navigate a series of dimensionally and mechanically defined environments to regenerate myofibers and replenish the stem cell pool (**Fig. 7**) (*15*, *21*). In homeostasis, MuSCs are positioned between a myofiber and the basal lamina; however, this structure is disrupted following muscle injury, prompting MuSCs to break their quiescence (*13*). After the removal of necrotic myofibers, MuSCs rapidly expand and differentiate on the remnant basal lamina (i.e., ghost fibers), where tissue stiffness decreases to approximately 5 kPa from the homeostatic 12 kPa (*14*, *15*, *21*). As regenerating myofibers re-emerge, MuSCs experience increasing confinement, with stiffness exceeding 25-30 kPa before returning to the homeostatic levels (*21*). Here, we model and investigate how these mechanical changes influence MuSC behavior by engineering 3D hydrogel bilayer platforms that mimic distinct stages of muscle regeneration, with variable confinement, and seeded primary MuSCs at the interface. Our findings show that 3D confinement regulates MuSC activation, proliferation, and differentiation through nuclear size and chromatin accessibility (**Fig. 7**). MuSCs in unconfined 2D conditions freely activate, expand, and exhibit large nuclei with high H4K16ac levels, while those in highly confined 3D environments show slower activation and smaller nuclei with reduced H4K16ac levels (**Fig. 7**). These results collectively reveal the critical role of physical confinement in slowing myogenic expansion and commitment through mechano-epigenetic mechanisms. Using the engineered *ex vivo* platform, we reveal that 3D mechanical cues are key modulators of MuSC behavior, providing a novel insight that is difficult to systematically study through conventional animal models.

**Figure 7.**
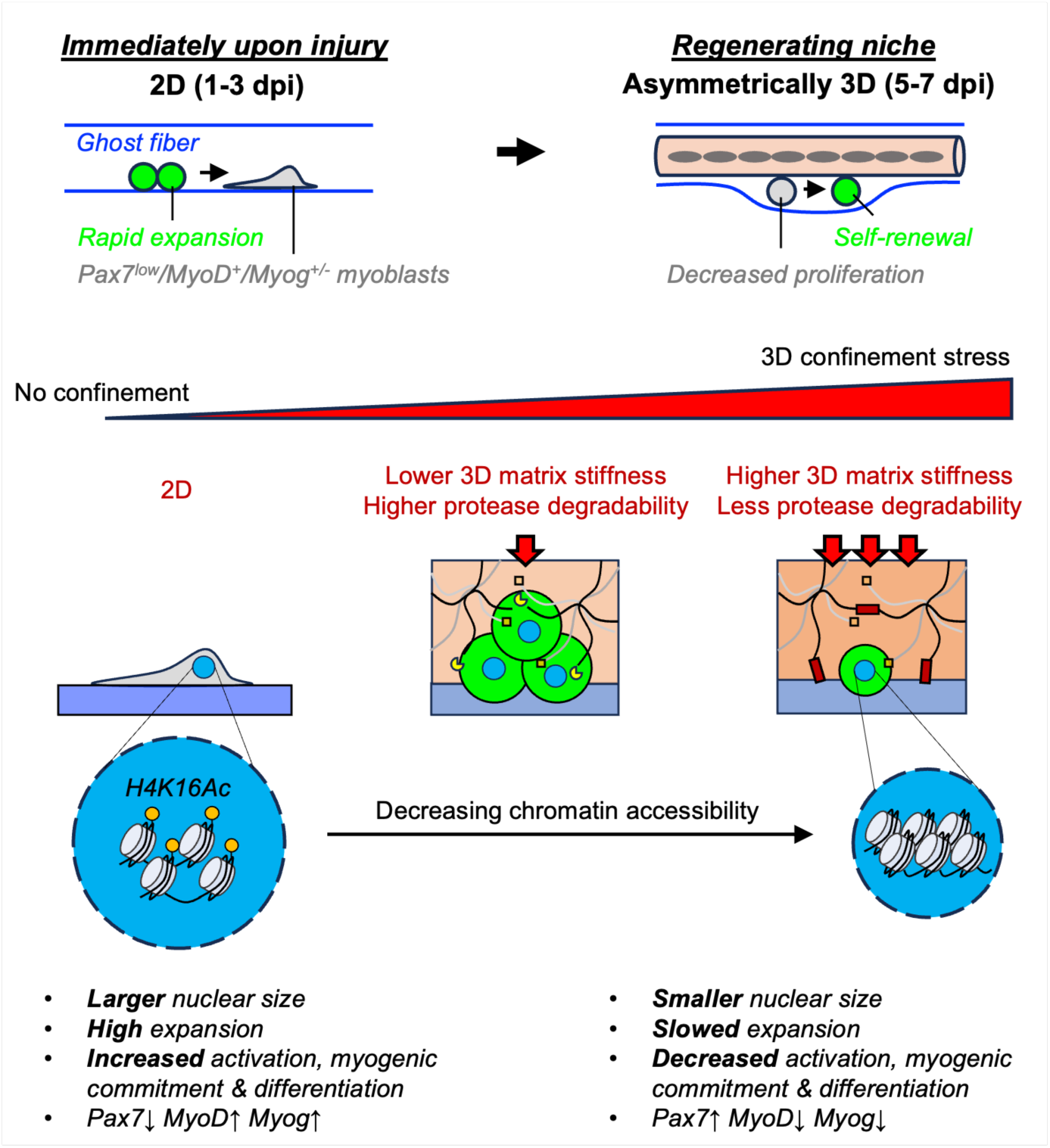
Proposed model of how confinement regulates MuSC function by modulating nuclear size and chromatin accessibility. *In vivo*, MuSCs initially expand and differentiate by using the ghost fibers as a 2D scaffold following injury. As regeneration progresses, MuSCs re-enter the 3D niche, where their proliferation decreases and self-renewal increases. It is unknown how mechanical confinement regulates these processes in muscle regeneration. In an unconfined 2D environment, MuSCs increase their nuclear size and acquire H4K16ac, which enhances their activation, proliferation, and myogenic progression. In contrast, as confinement increases, MuSCs exhibit reduced nuclear size and lower levels of H4K16ac, resulting in delayed activation, slower proliferation, and maintenance of a more stem-like state. We propose that confinement functions as a mechanical brake to inhibit persistent myogenic commitment through modulation of nuclear size and chromatin accessibility.

We demonstrated that exposure to physical confinement significantly decreases MuSC proliferative potential while maintaining *Pax7* expression and delaying MyoD and MYOG expression, effectively acting as a mechanical brake that inhibits differentiation compared to unconfined 2D conditions. These findings highlight the profound influence of the cell-extrinsic microenvironment on MuSC function. Consistent with our findings, a recent *in vivo* study showed that transplanting freshly isolated MuSCs either concurrently with injury or 5 days post-injury (dpi) resulted in a greater proportion of donor cells engrafting as MuSCs when transplantation occurred at 5 dpi, compared to those transplanted immediately after injury, which were more likely to differentiate and fuse (*16*). Similarly, in another 3D culture study, MuSCs introduced to 3D myotube templates exhibited robust engraftment as quiescent MuSCs with long cytoplasmic projections and polarity, a behavior not observed in 2D cultures (*34*, *45*). These observations suggest that the 3D mechanical and environmental cues present at later stages of regeneration, when the niche is more confining and mature, act as a brake that prevents uncontrolled expansion and differentiation, and signals MuSCs self-renewal. Together, these findings highlight the importance of confinement emergence in regulating MuSC behavior during muscle regeneration.

We observed that MuSCs maintained within the 3D bilayer hydrogels, particularly in the stiffest and most confining 35 kPa/12 kPa condition – mimicking the late stage of regeneration when stiffness exceeds homeostatic levels (*21*) – exhibited a further decrease in proliferation (**Fig. 2B, C**) and MyoD expression by day 5 (**Fig. 3C, D**). Nuclear size and the number of nuclei per colony were also significantly lower compared to other 3D bilayer hydrogel conditions (**Fig. 5A-F**), suggesting that MuSC function is regulated in a confinement-dependent manner. These findings also indicate that MuSC behavior is modulated progressively across a spectrum, with increasing confinement tuning cell fate decisions. Our findings align with a study demonstrating that apical compression promotes the return of MuSCs to quiescence (*24*). In that study, ex vivo compression was applied to induce deformation of MuSCs (*24*). However, our 35 kPa/12 kPa condition is less drastic and lacks the direct compressive force applied to the cells in the other model. This absence of compression likely explains why we did not observe an increase in Pax7EGFP intensity over time (**Fig. 2D**), suggesting that these MuSCs are not fully returning to quiescence under this level of confining stress alone in our system. It is also possible that a subset of proliferative MyoD+ MuSCs in the 35 kPa/12 kPa conditions undergo mitotic catastrophe, potentially due to mitotic compression that buckles the microtubule spindle (*46–50*), leading to their selective elimination and a slow decrease in cell number (**Fig. 2B**), ultimately favoring the survival of more quiescent MyoD-MuSCs (**Fig. 3C, D**).

This study demonstrated that MuSC nuclear size is regulated by the degree of confinement, with MuSCs on unconfined 2D substrates exhibiting the largest nuclei (**Fig. 5A-F**). Quiescent MuSCs are typically small, while Galert and activated MuSCs become larger (*15*, *34*, *51*), suggesting that MuSCs need space to expand, and that confinement may limit this process. Our results show that mechanical cues directly regulate nuclear size. In corroboration with our findings, studies have shown that the nucleus can transduce mechanical and spatial constraint cues into biochemical signals, effectively acting as a gauge of cellular deformation (*52*, *53*). In neural stem cells, confining stress similarly reduces nuclear size and alters chromatin architecture (*9*, *54*), supporting this notion.

Nuclear shape and deformation also directly impact chromatin architecture (*55–57*). In MuSCs, quiescence is associated with condensed chromatin and low transcriptional activity (*41*). Upon activation, MuSCs undergo epigenetic changes, including chromatin reorganization and histone modifications (*58*). In this study, we evaluated H4K16ac, (**Fig. 6A, B**) where lysine 16 acetylation in histone H4 relaxes chromatin and increases accessibility to the transcriptional machinery of myogenic genes (*42–44*). We observed that H4K16ac expression was positively associated with nuclear size, with more activated and larger MuSC nuclei on 2D and soft 3D bilayer hydrogels expressing higher levels of H4K16ac (**Fig. 6A, B**). Our findings suggest that changes in nuclear size due to variable confinement influence the epigenetic landscape of MuSCs, linking mechanical cues to potential changes in gene expression.

This study provides fresh insights into the role of 3D confinement in regulating MuSC behavior; however, it has a few limitations. First, while the engineered 3D bilayer hydrogel platform mimics certain aspects of the MuSC niche, it does not fully recapitulate the complexity of the *in vivo* microenvironment. Moreover, our system does not incorporate dynamic mechanical forces such as compression or tension, which likely play critical roles *in vivo*. Future studies incorporating dynamic mechanical forces, such as cyclic compression or tension, into 3D platforms could further elucidate how physical forces regulate MuSC behavior. The focus on H4K16ac as an epigenetic marker provides a partial and indirect view of the chromatin landscape, and further studies to include other epigenetic modifications may provide a more complete epigenetic profile that contributes to MuSC regulation under mechanical confinement. Nonetheless, the model presented here provides a tunable foundation for incorporating additional niche-derived factors and cells to better mimic the MuSC microenvironment in health, disease, and aging – offering a platform that enables systematic investigation of the physicochemical properties influencing MuSC function in steady-state, injury, and disease conditions, as well as a platform for cell manufacturing and regenerative therapies for skeletal muscle disorders.

## MATERIALS AND METHODS

### Mice

Animals were housed and bred in the Center for Comparative Medicine and Surgery Facility of the Icahn School of Medicine at Mount Sinai under the approved protocol by the Icahn School of Medicine at Mount Sinai Institutional Animal Care and Use Committee. C57BL6/J mice were acquired from the Jackson Laboratory (#000664). Pax7EGFP mice were provided by F. Mourkioti. Generation of Pax7EGFP has been published (*32*). Heterozygous Pax7EGFP was maintained by crossing with C57BL6/J wild-type mice. Genotyping was performed through Transnetyx.

### MuSC Isolation and Culture

MuSCs were isolated from male and female C57BL6/J or heterozygous Pax7EGFP mice, aged 4-8 weeks. MuSCs were isolated by magnetic-activated cell sorting (MACS), as described previously (*45*, *59*, *60*). Mouse muscle tissues were harvested from hindlimbs and minced using surgical scissors. The minced tissues were incubated in digest solution (2.5 U/ml Dispase II, MilliporeSigma and 0.2 w/v % Collagenase Type II, Worthington in DMEM) at 37°C for 90 minutes. Ham’s F-10 media containing 20% fetal bovine serum (FBS) was added to dilute the digest solution and filtered through a 0.70 µm cell strainer. Cells were washed through centrifugation (300g, 5 min, 4°C), then filtered again through a 0.35 µm cell strainer. Cells were incubated with biotinylated anti-mouse antibodies (CD31, 1:150, BioLegend; CD45, 1:150, BioLegend; and Sca-1, 1:150, BioLegend) at 4°C for 40 min. Cells were washed through centrifugation and labeled with 20 µL streptavidin beads (1:20) at 4°C for 20 min. Labeled cells were removed through negative selection using an LD column (Miltenyi). To further increase the purity of MuSCs, cells were incubated with anti-integrin ⍺-7 microbeads (1:12.5, Miltenyi) and positively selected using an LS column (Miltenyi). The enriched MuSCs were washed and filtered through a 35 µm cell strainer before use.

### MuSC Culture on Tissue Culture Plastic

µ-slides 8 well (Ibidi USA, #80821) were coated with coating solution (Collage, Type I, 5 ug/mL Gibco; Laminin, 10 ug/mL, Gibco) overnight in 4C. The coated slides were washed with phosphate-buffered saline (PBS) and dried completely before seeding the isolated MuSCs. 10,000 freshly isolated MuSCs were seeded per well. MuSCs were treated with 50 ng/mL of bFGF daily for the first 2 days in the growth media (Ham’s F-10, Gibco; 20% FBS, GeminiBio; 1X Penicillin-Streptomycin, Gibco; 1X GlutaMax; Gibco). Cultures were maintained at 5% CO2 and 37°C.

### Hydrogel Synthesis, Cell Encapsulation, and Culture

In this study, we synthesized poly(ethylene) glyocol norbornene (PEG-NB) hydrogels of varying Young’s modulus: 5 kPa (4% 20 kDa 4-arm PEG-NB, Creative PEGworks; 2 mM RGD, Genscript), 12 kPa (5% 40 kDa 8-arm PEG-NB, Creative PEGworks; 2 mM RGD, Genscript), and 35 kPa (7% 20 kDa 8-arm PEG-NB, Creative PEGworks; 2 mM RGD, Genscript). Flat bottom hydrogels were fabricated by first plasma treating µ-slides 8 well (Ibidi USA, #80821) to make the wells hydrophilic. 65 uL of the unpolymerized hydrogel solution (PEG-NB macromers; 2 mM RGD; 2 mM Lithium phenyl-2,4,6-trimethylbenzoylphospinate LAP, MilliporeSigma; 3.4 kDa PEG-dithiol crosslinker, MilliporeSigma) was added and a sterilized round 8-mm glass coverslip was gently placed on top of the solution at the center of the well. Hydrogel polymerized with 10 mW/cm^2^ 365 nm UV light for 2 minutes. The coverslips were removed, and hydrogels were swollen in PBS overnight at 4°C before cell seeding. After allowing cell attachment overnight, media from the well was gently removed and 65 uL of unpolymerized top hydrogel solution (PEG-NB macromers; 2 mM RGD; 1.2 mM LAP; VPM peptide crosslinker, Genscript) was added. After 45 seconds, the top hydrogel was polymerized with 10 mW/cm^2^ 365 nm UV light for 1 minute. The cells were cultured in growth media containing 50 ng/mL of bFGF and maintained at 5% CO2 and 37°C. C2C12 cell line was used for initial platform development and characterization.

### Hydrogel Compression Test

Mechanical compression test was performed to assess the Young’s modulus of the hydrogels. Hydrogel samples were fabricated using a custom mold with a diameter of 4 mm and a height of 1.7 mm. The samples were incubated in PBS at 4°C overnight to allow for swelling. After this period, the diameter and thickness of the hydrogels were measured. The samples were then subjected to displacement-controlled unconfined compression using the Electroforce 3200 testing instrument (TA Instruments, New Castle, DE) in conjunction with WinTest 7 software (TA Instruments) at a strain rate of 0.2% per second, compressing until a total strain of 50% was achieved. Young’s compressive modulus was determined by calculating the slope of the first linear region before 25% strain (*61*).

### Pax7EGFP Live Cell Imaging and Quantification

After seeding MuSCs onto 2D 12 kPa hydrogels and in 3D hydrogels (comprising top layers of 5/12 kPa, 12/12 kPa, and 35/12 kPa), the samples were imaged in vitro using a Leica DMi8 THUNDER 3D Imager equipped with an LED5 fluorescence light source and HC PL Fluotar 20X/0.55 objective. EGFP imaging was conducted with consistent exposure and intensity settings across time points (488 nm wavelength, power at 8%, and a 20X objective). Each experimental group consisted of four hydrogels, and three regions per well were captured for analysis. The EGFP intensity was quantified by outlining cells and measuring the grayscale intensity using ImageJ/FIJI, which also enabled the scoring of cell numbers based on EGFP fluorescence. Normalized mean Pax7EGFP intensity is expressed as 100% at the mean of all groups at 2h and 0% is set to the lowest single value at 120h. The number of colonies per image was analyzed manually.

### Immunofluorescence

The samples were fixed with 4% paraformaldehyde (PFA) for 30 minutes. The samples were digested with 100 U/mL collagenase at 37°C for 3 hours to degrade the top hydrogel layer. Following digestion, the cells were incubated with a blocking/permeabilization buffer (5% goat serum, 2% bovine serum albumin, 0.5% Triton X-100 in PBS) at room temperature for one hour. The samples were then treated with primary antibodies [Anti-PAX7, 1:200, DSHB; Anti-MyoD (G-1), 1:500, Santa-Cruz; Anti-Myogenin (F5D), 1:500, Santa-Cruz; Anti-Acetyl-Histone H4 (Lys16) (E2B8W), 1:500, Cell Signaling Technology] overnight at 4°C. The samples were then washed with PBS-T (0.05% Tween-20 in PBS) and incubated with secondary antibodies [Alexa Fluor 546 Goat anti-Mouse, 1:1000, Invitrogen; Alexa Fluor Plus 647 Goat anti-Rabbit, 1:1000, Invitrogen] and Hoechst (1:1000, ThermoFisher Scientific) for 3 hours at room temperature. After incubation, the samples were washed overnight with PBS-T, which included at least five changes of fresh washing buffer. Immunofluorescence imaging was performed using a Leica DMi8 THUNDER 3D Imager equipped with HC PL Fluotar 20X/0.55 objective and HCX PL Fluotar L 40X PH2 objective.

### Image Analyses

The percentage of positivity of MyoD, MYOG, H4K16ac, and PAX7 was calculated by manually counting the labeled nuclei relative to the total Hoechst+ nuclei using ImageJ/FIJI. Additionally, nuclear size was quantified by outlining and measuring Hoechst-stained nuclei using ImageJ/FIJI.

### Statistical Analyses

Statistical analyses were performed using GraphPad Prism. Two-tailed t-test, one-way analysis of various (ANOVA) with Tukey’s post-hoc analysis, two-way ANOVA with Dunn’s post-hoc analysis, two-tailed Mann-Whitney U test, and Kruskal Wallis with Dunn’s post-hoc analysis was performed based on data normality and the number of comparisons. Statistical significance was set at p<0.05.

## ACKNOWLEDGMENTS/FUNDING

We are deeply grateful to Dr. Robert Krauss for his insightful discussions and careful reading of the manuscript. Research reported in this publication was supported by the National Institute of Arthritis and Musculoskeletal And Skin Diseases of the National Institutes of Health under Award number R01AR080616 to W.M.H. The content is solely the responsibility of the authors and does not necessarily represent the official views of the National Institutes of Health.

## AUTHOR CONTRIBUTIONS

G.P. and J.A.G. performed experiments and data collection. F.M. provided animals. G.P., J.A.G., W.M.H. conceptualized, designed the studies, and analyzed the data. G.P., J.A.G., F.M., and W.M.H. interpreted data and wrote the manuscript.

## DATA AND MATERIALS AVAILABILITY

All data needed to evaluate the conclusions in the paper are present in the paper and the Supplementary Materials.

---

**Supplemental Figure 1.**
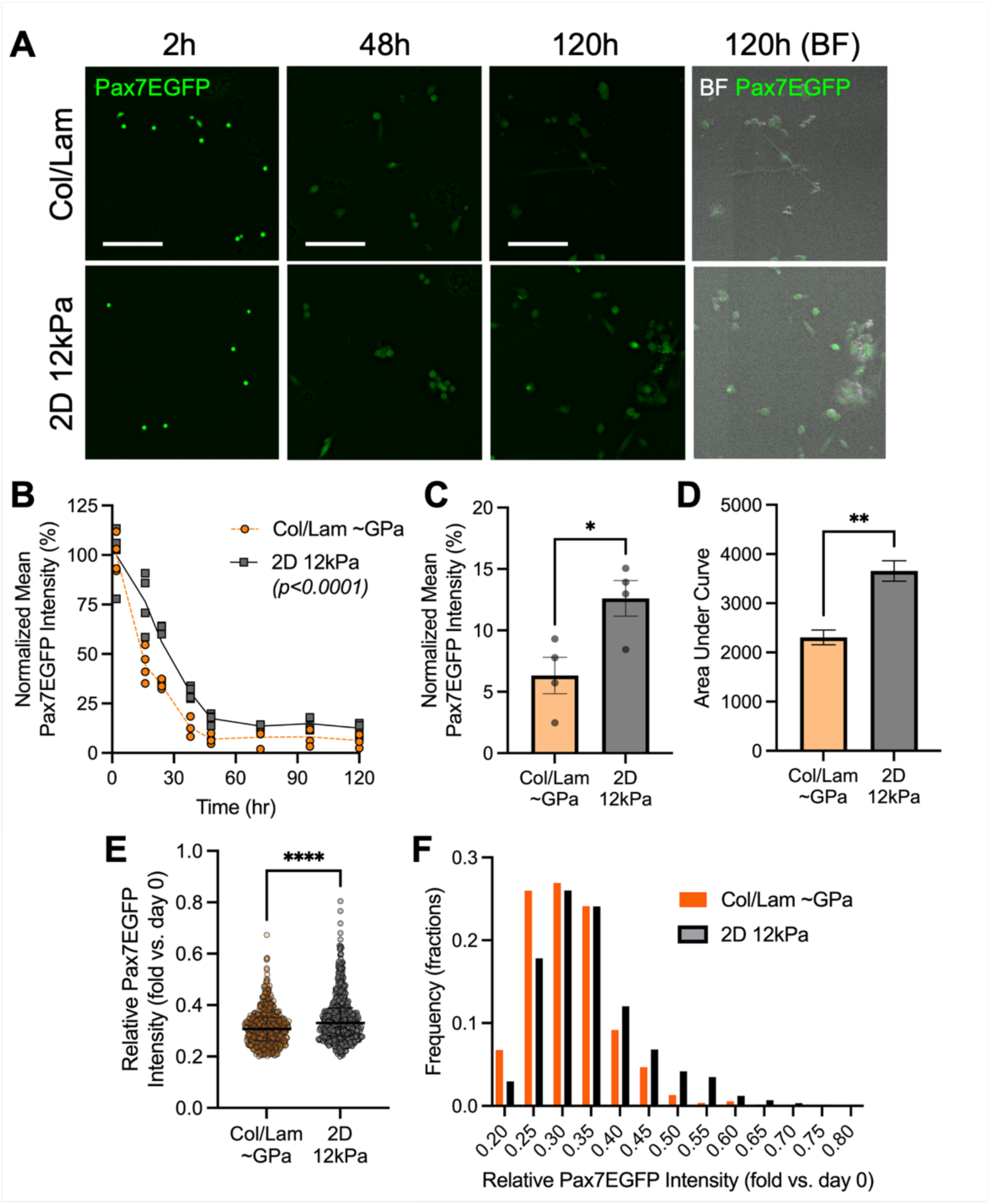
Dynamic Pax7EGFP reporter MuSCs enable longitudinal tracking of Pax7 expression ex vivo. **(A)** Representative micrographs of Pax7EGFP MuSCs cultured on collagen/laminin (col/lam)-coated tissue culture plastic and RGD-functionalized 12 kPa hydrogel. Scale bar: 100 µm. BF: Bright Field. **(B)** Normalized mean Pax7EGFP intensity over time. p<0.0001 via 2-way ANOVA. **(C)** Normalized mean Pax7EGFP intensity at 120-hour. n=4 hydrogels. * p<0.05 via two-tailed t-test. **(D)** Area under curve analyses of (B). ** p<0.01 via two-tailed t-test. **(E)** Relative Pax7EGFP intensity of MuSCs on day 5 normalized to day 0 intensity. n=535-573 cells analyzed. **** p<0.0001 via two-tailed Mann-Whitney U test. **(F)** Frequency distribution of relative Pax7EGFP intensity of MuSCs normalized to day 0 intensity.

**Supplemental Figure 2.**
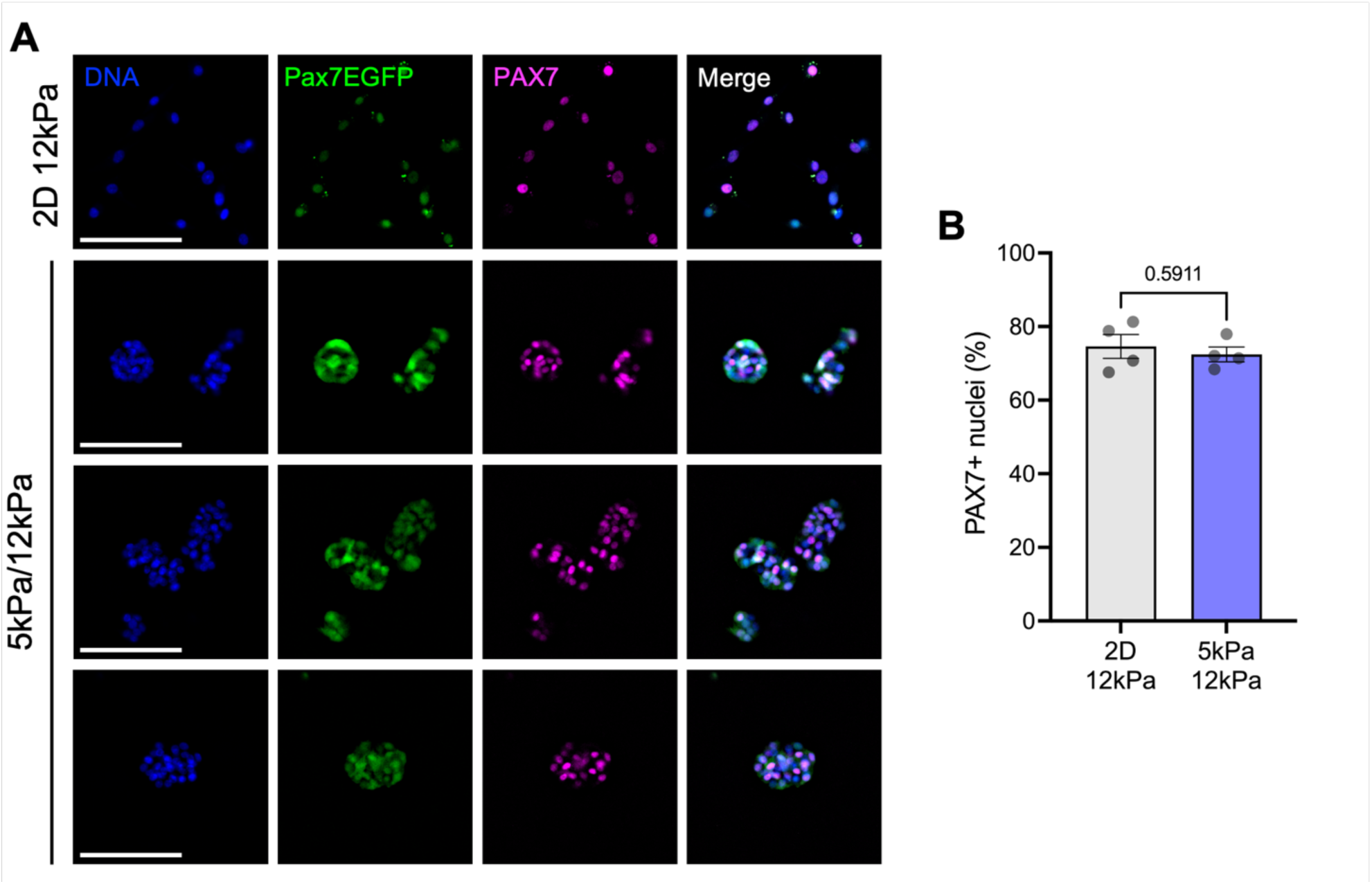
Percent PAX7 positivity is unaffected by confinement by day 5. **(A)** PAX7 immunostaining on day 5. Scale bar: 100 µm. **(B)** Percentage of PAX7+ nuclei on day 5. n=4 hydrogels. p=0.5911 via two-tailed unpaired t-test.

**Supplemental Figure 3.**
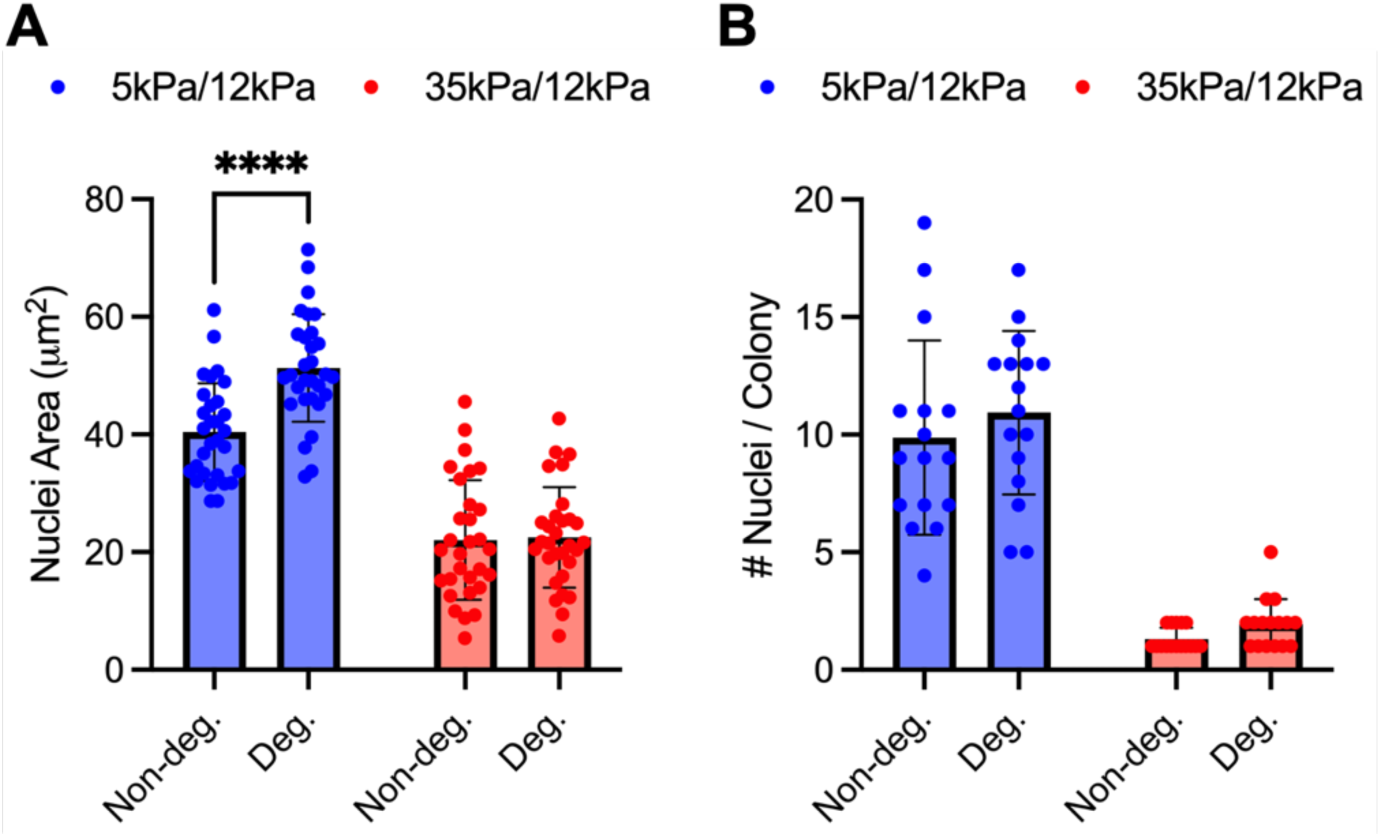
MuSC confinement is primarily regulated by stiffness rather than protease-mediated hydrogel degradation. **(A)** Quantification of nuclei area. Day 5. **** p<0.0001 via two-way ANOVA with Dunn’s post-hoc. n=30 nuclei analyzed. **(B)** Quantification of number of nuclei per colony. Day 5. n=16 colonies analyzed.

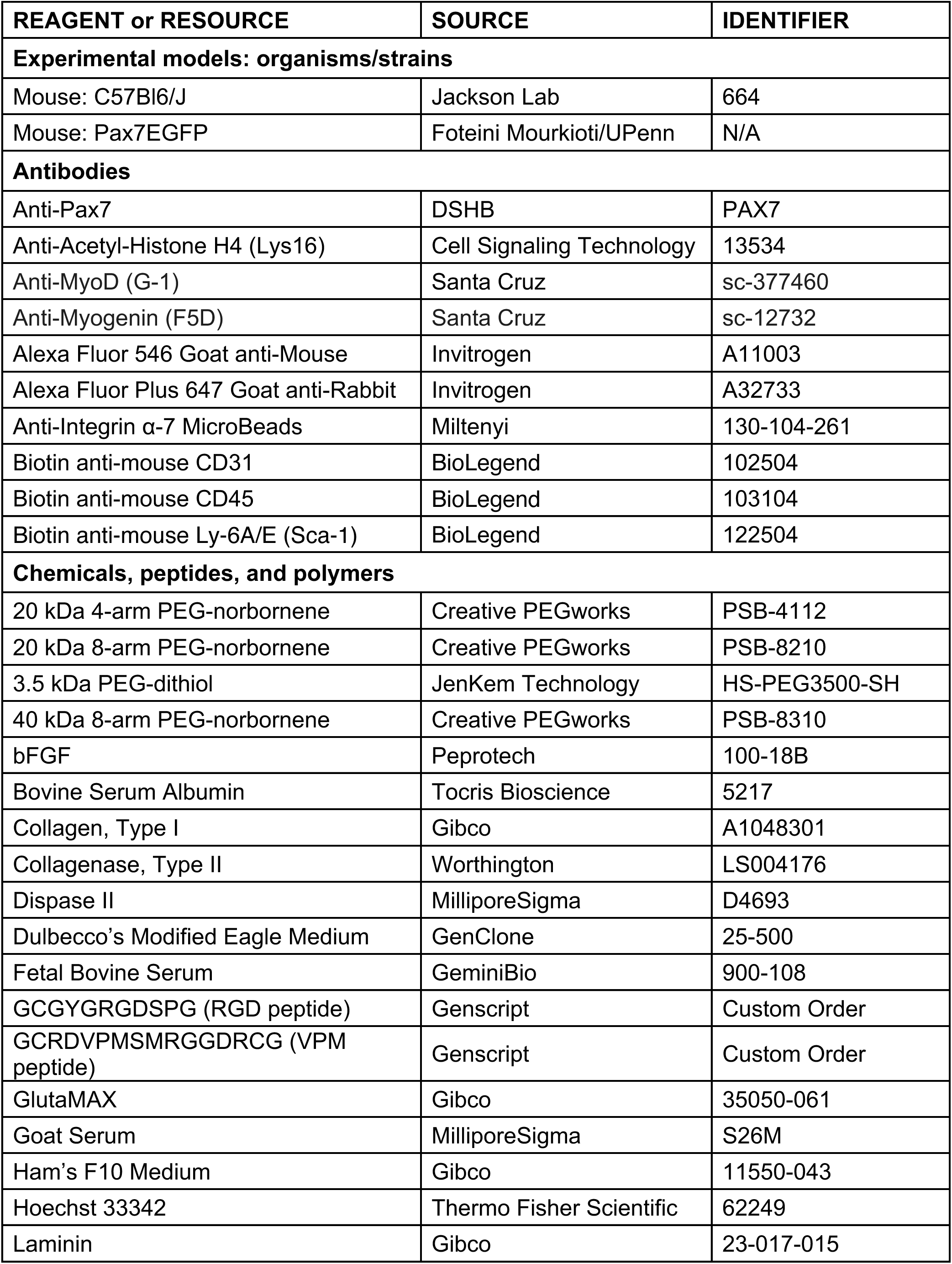
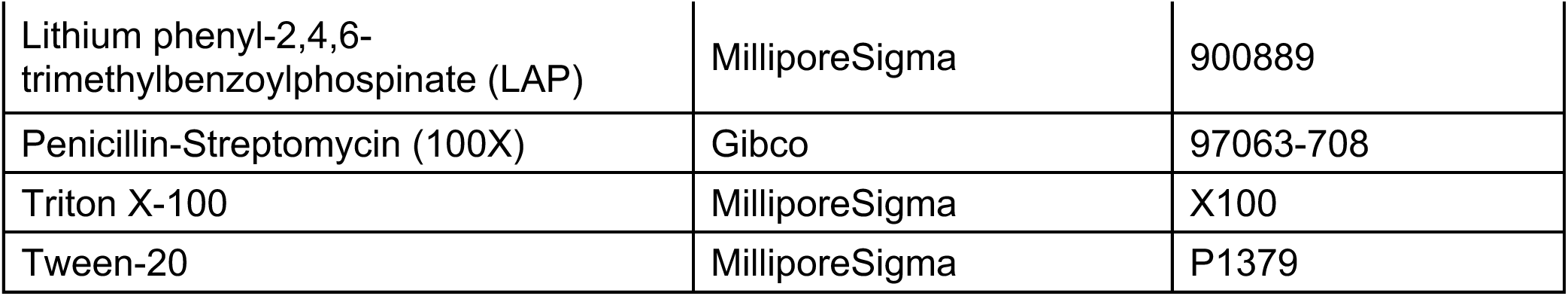
Summary Table of Reagent & Resources.

