## Supplementary Materials for "3D mechanical confinement directs muscle stem cell fate and function"

Woojin M. Han, PhD
Assistant Professor
Department of Orthopaedics
Department of Cell, Developmental, and Regenerative Biology
Institute for Regenerative Medicine
Black Family Stem Cell Institute
Icahn School of Medicine at Mount Sinai
Annenberg Building, Floor 20-66A, Box 1188
1468 Madison Ave, New York, NY 10029

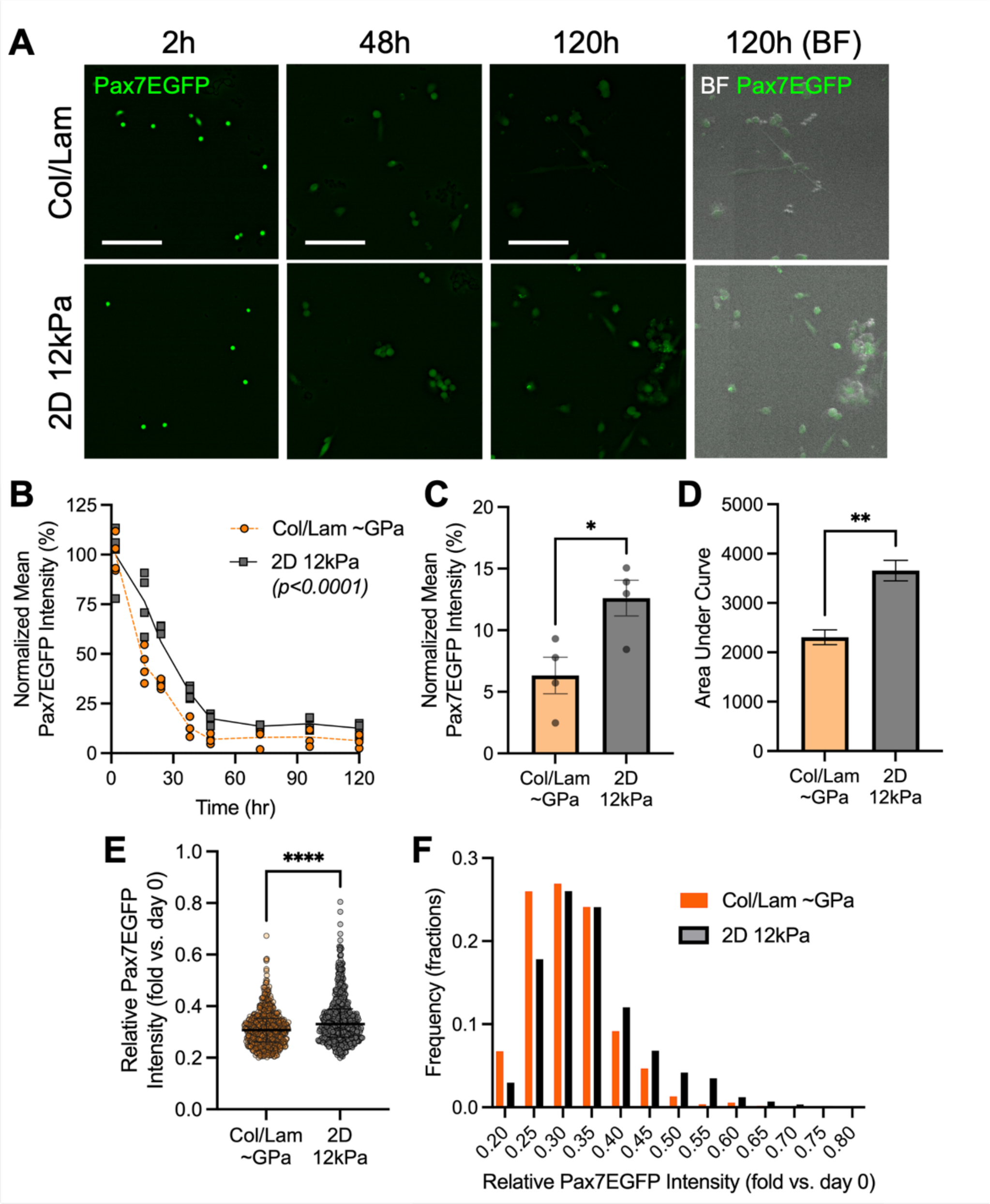


**Supplemental Figure 1. Dynamic Pax7EGFP reporter MuSCs enable longitudinal tracking of Pax7 expression ex vivo. (A)** Representative micrographs of Pax7EGFP MuSCs cultured on collagen/laminin (col/lam)-coated tissue culture plastic and RGD-functionalized 12 kPa hydrogel. Scale bar: 100 μm. BF: Bright Field. **(B)** Normalized mean Pax7EGFP intensity over time. $p<0.0001$ via 2-way ANOVA. **(C)** Normalized mean Pax7EGFP intensity at 120-hour. n=4 hydrogels. * $p<0.05$ via two-tailed t-test. **(D)** Area under curve analyses of (B). ** $p<0.01$ via two-tailed t-test. **(E)** Relative Pax7EGFP intensity of MuSCs on day 5 normalized to day 0 intensity. n=535-573 cells analyzed. **** $p<0.0001$ via two-tailed Mann-Whitney U test. **(F)** Frequency distribution of relative Pax7EGFP intensity of MuSCs normalized to day 0 intensity.

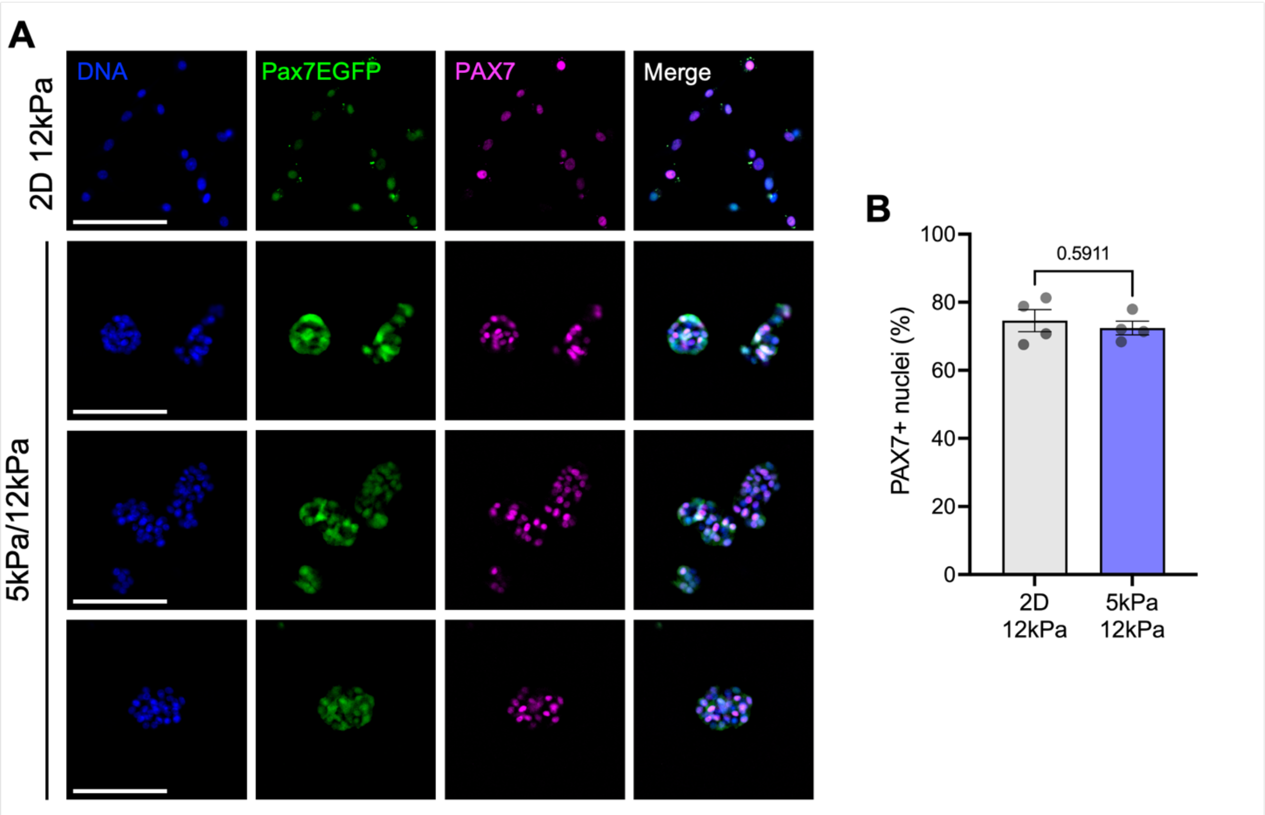


**Supplemental Figure 2. Percent PAX7 positivity is unaffected by confinement by day 5. (A)** PAX7 immunostaining on day 5. Scale bar: 100 µm. **(B)** Percentage of PAX7+ nuclei on day 5. n=4 hydrogels. p=0.5911 via two-tailed unpaired t-test.

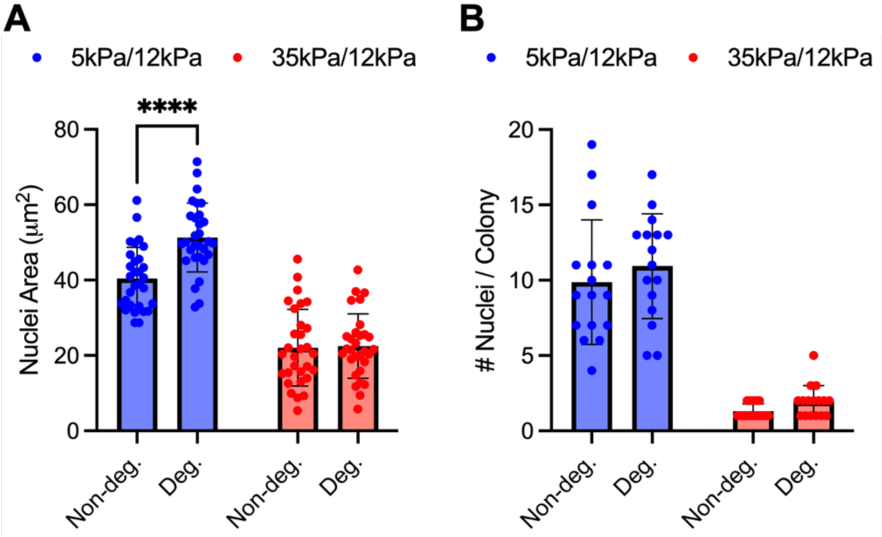


**Supplemental Figure 3. MuSC confinement is primarily regulated by stiffness rather than protease-mediated hydrogel degradation. (A)** Quantification of nuclei area. Day 5. **** p<0.0001 via two-way ANOVA with Dunn's post-hoc. n=30 nuclei analyzed. **(B)** Quantification of number of nuclei per colony. Day 5. n=16 colonies analyzed.

**Summary Table of Reagent & Resources**

| REAGENT or RESOURCE | SOURCE | IDENTIFIER |
|---|---|---|
| **Experimental models: organisms/strains** | | |
| Mouse: C57Bl6/J | Jackson Lab | 664 |
| Mouse: Pax7EGFP | Foteini Mourkioti/UPenn | N/A |
| **Antibodies** | | |
| Anti-Pax7 | DSHB | PAX7 |
| Anti-Acetyl-Histone H4 (Lys16) | Cell Signaling Technology | 13534 |
| Anti-MyoD (G-1) | Santa Cruz | sc-377460 |
| Anti-Myogenin (F5D) | Santa Cruz | sc-12732 |
| Alexa Fluor 546 Goat anti-Mouse | Invitrogen | A11003 |
| Alexa Fluor Plus 647 Goat anti-Rabbit | Invitrogen | A32733 |
| Anti-Integrin α-7 MicroBeads | Miltenyi | 130-104-261 |
| Biotin anti-mouse CD31 | BioLegend | 102504 |
| Biotin anti-mouse CD45 | BioLegend | 103104 |
| Biotin anti-mouse Ly-6A/E (Sca-1) | BioLegend | 122504 |
| **Chemicals, peptides, and polymers** | | |
| 20 kDa 4-arm PEG-norbornene | Creative PEGworks | PSB-4112 |
| 20 kDa 8-arm PEG-norbornene | Creative PEGworks | PSB-8210 |
| 3.5 kDa PEG-dithiol | JenKem Technology | HS-PEG3500-SH |
| 40 kDa 8-arm PEG-norbornene | Creative PEGworks | PSB-8310 |
| bFGF | Peprotech | 100-18B |
| Bovine Serum Albumin | Tocris Bioscience | 5217 |
| Collagen, Type I | Gibco | A1048301 |
| Collagenase, Type II | Worthington | LS004176 |
| Dispase II | MilliporeSigma | D4693 |
| Dulbecco's Modified Eagle Medium | GenClone | 25-500 |
| Fetal Bovine Serum | GeminiBio | 900-108 |
| GCGYGRGDSPG (RGD peptide) | Genscript | Custom Order |
| GCRDVPMSMRGGDRCG (VPM peptide) | Genscript | Custom Order |
| GlutaMAX | Gibco | 35050-061 |
| Goat Serum | MilliporeSigma | S26M |
| Ham's F10 Medium | Gibco | 11550-043 |
| Hoechst 33342 | Thermo Fisher Scientific | 62249 |
| Laminin | Gibco | 23-017-015 |

| | | |
|---|---|---|
| Lithium phenyl-2,4,6-trimethylbenzoylphospinate (LAP) | MilliporeSigma | 900889 |
| Penicillin-Streptomycin (100X) | Gibco | 97063-708 |
| Triton X-100 | MilliporeSigma | X100 |
| Tween-20 | MilliporeSigma | P1379 |
